# Batch cultivation of non-transgenic males suitable for SIT programs

**DOI:** 10.1101/828822

**Authors:** Siba Das, Maciej Maselko, Ambuj Upadhyay, Michael J. Smanski

## Abstract

The field performance of Sterile Insect Technique (SIT) is improved by sex-sorting and releasing only the sterile males. This can be accomplished by resource-intensive separation of males from females by morphology. Alternatively, sex-ratio biasing genetic constructs can be used to selectively kill one sex without the need for manual or automated sorting, but the resulting genetically engineered (GE) control agents would be subject to additional governmental regulation. Here we describe and demonstrate a method for the batch production of non-GE males that is applicable for sex-selective production of males suitable for genetic biocontrol programs. This method could be applied to generate the heterogametic sex (XY, or WZ) in any organism with chromosomal sex determination. We observed up to 100% sex-selection with batch cultures of more than 10^3^ individuals. Using a stringent transgene detection assay, we demonstrate the potential of mass rearing of transgene free males.

## Introduction

Insect pests impose a major burden to food production and human health worldwide. The most successful population control method in use today is the sterile insect technique (SIT)^1^. SIT relies on mass rearing of pest insects followed by a sterilization treatment (*e.g*. X-ray irradiation). Sterilized insects are released into the wild where sterile males compete with wild males to mate with wild females. Since females of many pest insects only mate once in their lifetime, mating with a sterile male prevents successful reproduction. Sufficiently large releases of sterile insects can be used to eliminate wild populations or prevent their establishment in a new area^2^. Also, SIT is considered safe to humans and the environment, as there are less off-target impacts compared to the application of chemical pesticides.

Existing SIT programs are used to control several major agricultural pests including the New World Screwworm (*Cochliomyia hominivorax*)^3^, Mediterranean Fruit Fly (*Ceratitis capitata*)^4^, and Queensland Fruit Fly (*Bactrocera tryoni*)^5^. All together, these programs produce and release billions of sterile insects on a weekly basis. The application of SIT to control the New World Screwworm resulted in the eradication of this livestock pest in North and Central America, with an estimated value of over $US 1 Billion/year^3^.

SIT for many insects, including *C. hominivorax* and *B. tryoni*, currently involves releasing both sterilized males and females. However, the effectiveness of SIT can be substantially increased if only males are released since they will then seek out wild females instead of mating with co-released sterile females^6^. In some insect pests, such as the Yellow Fever Mosquito (*Aedes aegyptiI)* and Spotted Wing Drosophila (*Drosophila suzukii)*, it is crucial that SIT programs only release males since sterile females can vector disease or damage crops.

A variety of sex-sorting techniques have been developed. Mechanical separation of *Aedes aegypti* pupae based on size differences^7^ can be effective and flow cytometric separation of transgene expressing female *Anopholes gambiae* has been demonstrated^8^; however, these approaches can be labor-intensive or require sophisticated equipment. Combining irradiation with heat-shock is female-lethal in mutant strains of *C. capitata* and is presently in use^9^. Repressible transgenic female-elimination constructs act as genetic biocontrol systems on their own and have been developed for *Ae. aegypti*^10^*, C. hominivorax*^11^, Sheep Blow Fly (*Lucilia cuprina*)^12^, Diamondback moth (*Plutella xylostella*)^13^, Pink Bollworm (*Pectinophora gossypiella*)^13^, and Silkworm (*Bombyx mori*)^14^. Public resistance and regulatory hurdles have unfortunately limited the broad use of released transgenic insects for pest control despite their effectiveness, specificity, reduction of insecticide use, and safety.

We describe here Subtractive Transgene Sex Sorting (STSS) which is a genetic approach to produce non-transgenic males. STSS relies on two transgenic strains; each of which has a lethal genetic circuit that can be repressed (Fig. 1a). One of the strains has the lethal circuit on the Y-chromosome (YL strain) and the other has the lethal circuit on the X-chromosome (XL strain). Non-transgenic males are produced in a two-step mating scheme (Fig. 1b). First, the true-breeding YL strain is grown in media that activates the lethal circuit, resulting in non-transgenic females in the subsequent generation. These non-transgenic females are combined with the XL strain in media that activates the lethal circuit. Mating between the XL males and non-transgenic females results in non-transgenic males. All other offspring die from expression of the X-linked lethal circuit. This technique is transferable to any organism that relies on genetic, as opposed to environmental, sex determination^15^.

**Fig. 1.**
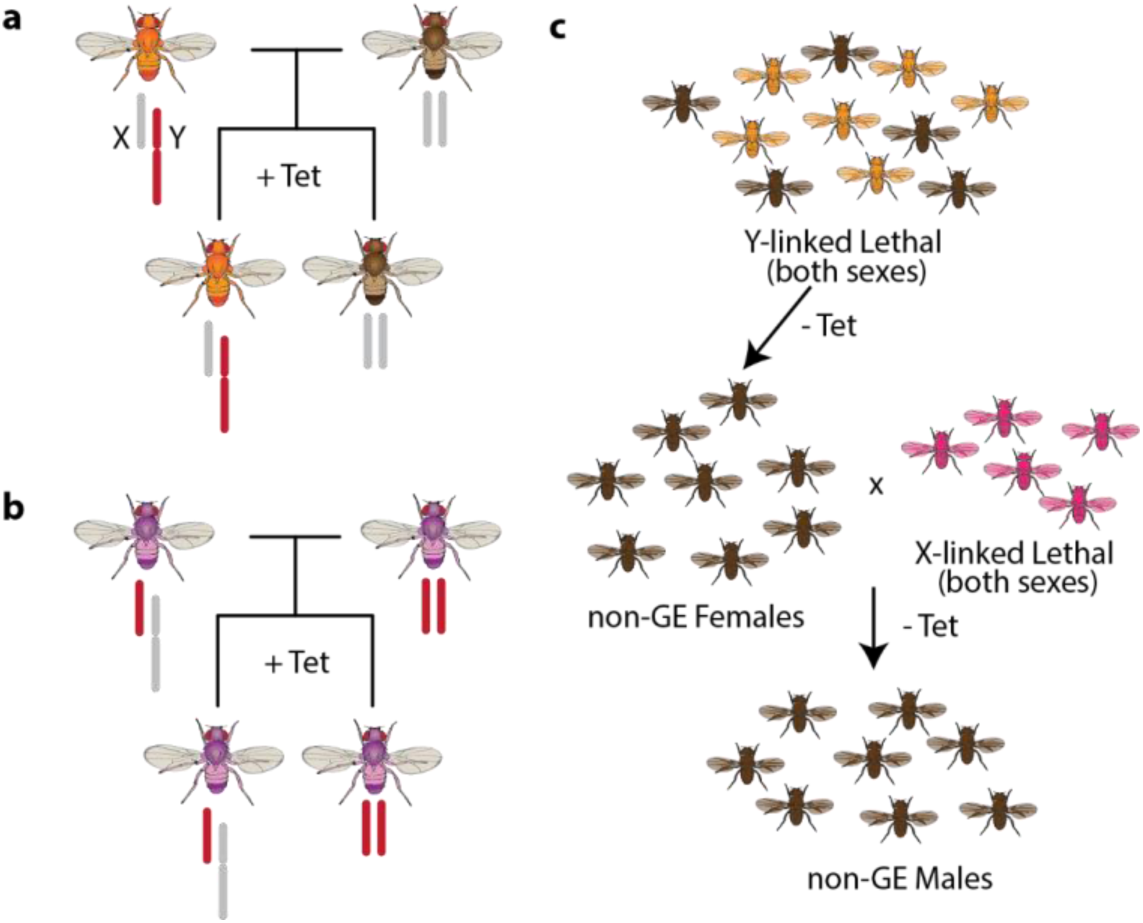
Overview of STSS. Minimal requirements for each strain to be used in STSS, including true-breeding population with conditional Y-linked lethality (**a**) or conditional X-linked lethality (**b**). (**c**) Mating scheme in absence of lethal gene repressor. Combining non-transgenic females produced from the YL strain with adult flies from the XL strain results in death of all offspring except for non-transgenic males.

## Results

### Design and construction of a repressible lethal transgenic construct

A repressible lethal genetic construct can be designed with a conditionally-expressed promoter driving a toxic gene product. We selected a tetracycline-repressible *hsp70* minimal promoter (*pHsp70*) due to its well-characterized behavior in model and applied insect species^12,16^. To drive lethality, we expressed the tet-transactivator (tTA), whose VP64 transactivation domain is toxic to cells when strongly expressed. We made tTA expression constructs with a positive feedback loop wherein *pHsp70* drives basal expression of tTA similar to what has been previously described^17,18^ (Fig. 2a). The presence of tetracycline prevents transactivation by tTA and keeps the construct expressed at basal, sub-lethal levels. In the absence of tetracycline, tTA binds to operators upstream of *pHsp70* and establishes a positive feedback loop which generates lethal amounts of tTA. We incorporated an MHC intron and syn21 5’-UTR translational enhancer features^19^ to further boost tTA expression. Tuning gene expression in lethal transgenic constructs is a balance between (i) incurring fitness effects from leaky expression in the repressed ‘off’-state and (ii) incomplete penetrance due to weak expression in the de-repressed ‘on’-state. Our priority was to ensure complete penetrance of the lethal phenotype in the derepressed state, so we designed the construct to favor strong expression.

**Fig. 2.**
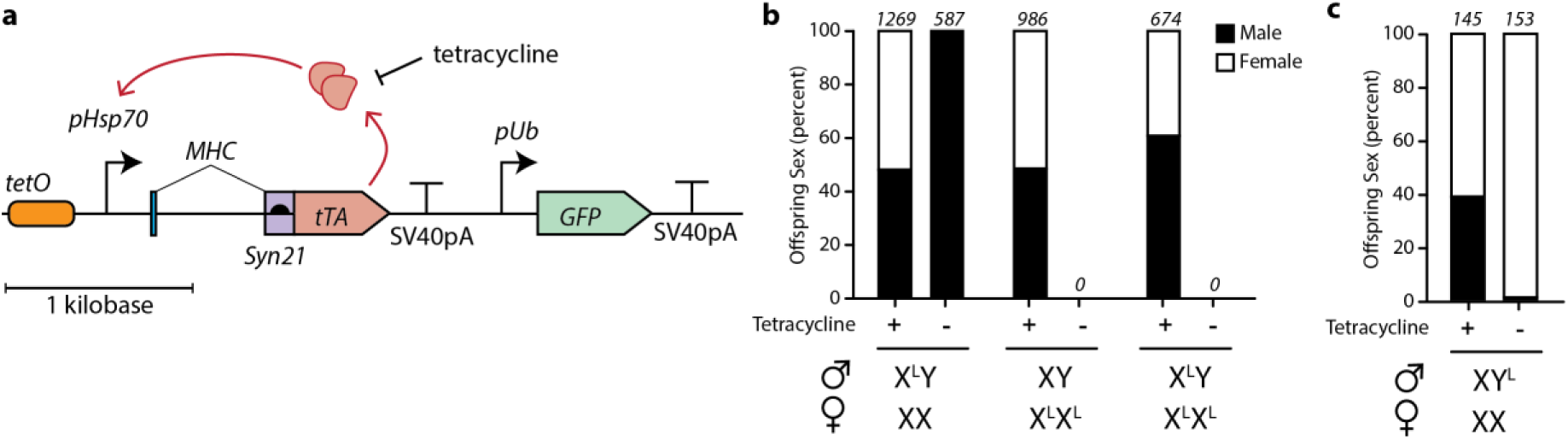
Sex-chromosome linked tet-repressible lethal circuits are effective. (**a**) Construct-level schematic of tet-repressible lethal circuit used in this study. (**b**) Proportion of male and female offspring generated from self-mating of DmXL^tTA^ or with non-transgenic (w1118) flies in presence or absence of tetracycline. Genotypes of parental flies are indicated below x-axis. Numbers above bars indicate total number of progeny produced from six biological replicates. (**c**) Proportion of male and females generated from mating DmYL^tTA^ with non-transgenic (w1118) flies in presence or absence of tetracycline. Numbers above bars show total number of progeny produced from three biological replicates.

To test the genetic circuit design, we engineered *D. melanogaster* due to its powerful genetic toolkit and also because its serves as a model for other insect pests. We used ΦC31-mediated transgenesis to integrate a single copy of the tTA circuit into *Att*P landing sites on the X and Y chromosomes of two separate strains (Fig. 2b, referred to as DmXL^tTA^ and DmYL^tTA^ from here on). Both of these strains were maintained in the presence of 100 µg/ml tetracycline. The genotype of transgenic flies was confirmed by PCR amplification (data not shown). The DmXL^tTA^ were mated with *FM6*, an X-chromosome balancer strain, and then selfed to screen for females homozygous for the modified X-chromosome, which was confirmed by PCR (data not shown). From this point on, DmXL^tTA^ and DmYL^tTA^ were maintained as true-breeding lines in the presence of tetracycline.

### Performance of repressible lethal genetic constructs

To test the efficiency of toxic gene expression, virgin females and males (3 each) were mated on media lacking tetracycline. In each of at least three replicate crosses, no DmXL^tTA^ offspring survived to adulthood (Fig. 2b). This suggests that the repressible lethal transgenic construct is sufficiently strong to cause lethality in two copies (females) or one copy (males). In an analogous experiment with DmYL^tTA^ flies, four replicate crosses produced a total of 204 females and only 1 male (99.5% female). This single male did not reproduce when subsequently mated with non-transgenic females. Thus, both DmXL^tTA^ and DmYL^tTA^ produced a sufficiently lethal phenotype in the absence of tetracycline to remove the transgene from the accessible gene pool (Fig. 2bc).

### Sub-stoichiometric ratio of mixed-sex DmXL^tTA^ to female DmYL^tTA^ sufficient for non-transgenic male production

Non-transgenic males can be generated by crossing non-transgenic females produced by the DmYL^tTA^ strain and males from a mixed-sex true-breeding population of DmXL^tTA^ flies (Fig. 1c). We hypothesized that the number of non-transgenic males produced will be directly related to the number of non-transgenic mothers, but robust to decreasing numbers of DmXL^tTA^ fathers. This would be important for economically scaling-up the production of non-trangenic males for SIT programs. We performed experimental crosses between non-transgenic females and DmXL^tTA^ mixed-sex populations to determine the minimum sufficient ratio of parental genotypes. We observed a monotonically increasing number of total offspring produced as the ratio of DmXL^tTA^ males to DmYL^tTA^ females increased from 1:20 to 3:10 (Fig. 3a). The offspring number appeared to plateau or even decline after further increasing the number of DmXL^tTA^ males. This suggests that a ~1:3 ratio is sufficient to ensure that the number of DmXL^tTA^ males are not limiting the total number of offspring produced.

**Fig. 3.**
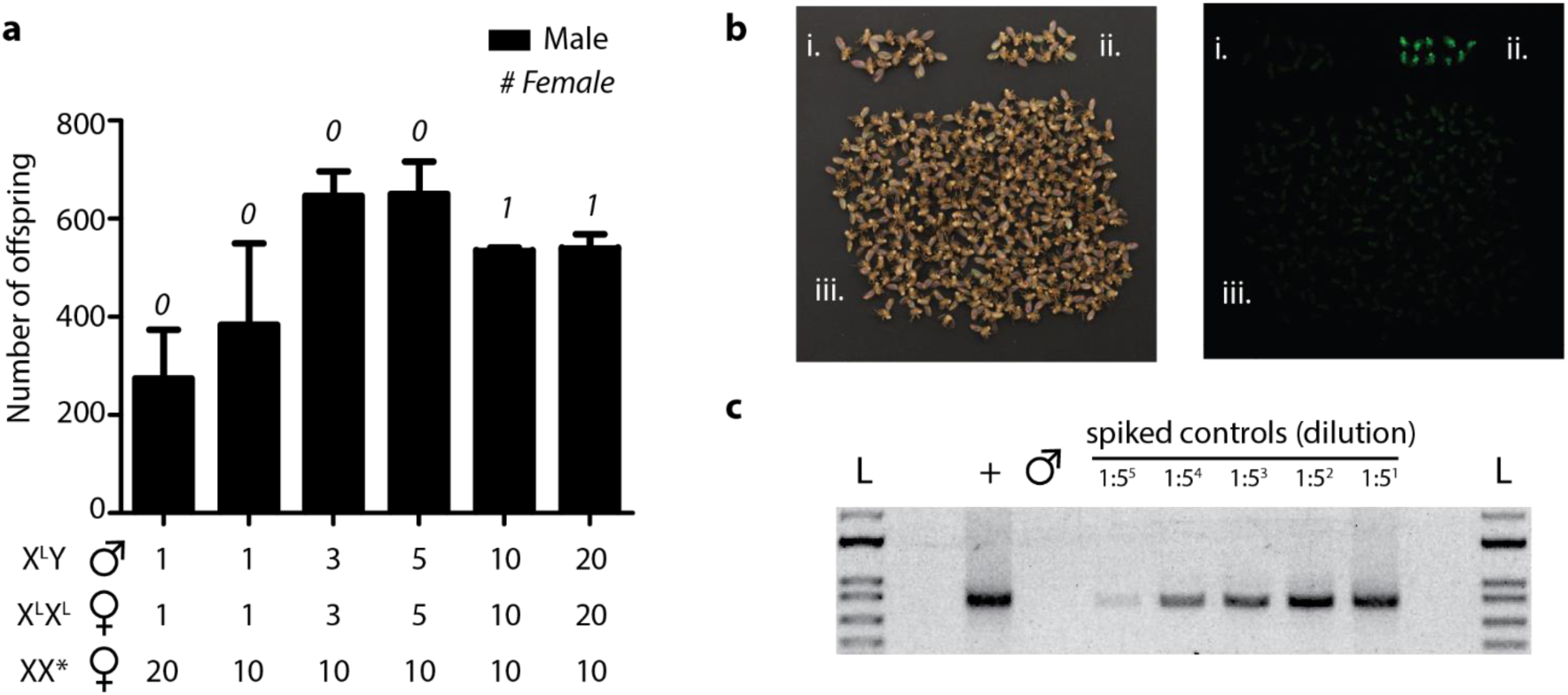
Batch production of adult males via STSS. (**a**) Average number of adult males obtained from mating between different proportions of non-transgenic female flies obtained from DmYL^tTA^ in absence of tetracycline (‘XX*’) when combined with adult DmXL^tTA^ flies in absence of tetracycline. Data represent mean numbers from 2-3 of biological replicates with error bars showing standard error of the mean (s.e.m.). Average numbers of females produced are indicated numerically above bars. Numbers below x-axis indicate ratio of genotypes in parental generation. Total numbers from all replicates are (from left to right): N=822, N=1151, N=1292, N=1301, N=1068, N=1078 (**b**) Bright-field (left) and fluorescent (right) images of parental (**i**) 10 DmYL^tTA^ females produced in the absence of tetracycline (‘XX*’), parental (ii) 10 DmXL^tTA^ males, and (iii) approximately 400 offspring from the batch production of non-transgenic males. Flies in (i) and (ii) are only included for visual comparison to the STSS males; they were not present in the final batch of STSS flies. (**c**) PCR amplification of transgene cassette from genomic DNA isolated from 10 DmXL^tTA^ flies (+), 1180 batch-produced STSS males (male symbol), or STSS gDNA spiked with DNA from DmXL^tTA^ flies at 5-fold dilutions from 1:5 (right) to 1:3125 (left). L denotes 1 kb plus DNA ladder.

At or below the optimal ratios of DmXL^tTA^ males to DmYL^tTA^ females, we observed 100% male offspring (N_combined_ = 5388 male offspring, 0 female offspring). We observed a total of 4 female offspring across all replicates when the ratio of DmXL^tTA^ males to DmYL^tTA^ females was 10:10 or 20:10 (N_combined_ = 2142 male offspring, 4 female offspring). It is unclear how these females were able to survive, but they lacked a GFP phenotype and thus did not appear to be transgenic (data not shown).

### Large-scale cultivation of non-transgenic males suitable for egg release

Next we tested the effectiveness of producing non-transgenic males by batch cultivation, following the mating scheme in Fig. 1c. We transferred a true-breeding culture of DmYL^tTA^ to media lacking tetracycline and cleared all of the adults from bottles after 5 days. Approximately 40 offspring emerged on the tetracycline-free medium. Adults from a true-breeding population of DmXL^tTA^ flies were added at the sub-stoichiometric ratio of 1:2. From this mating 1792 males and one female emerged. None of the more than 1000 males screened contained the GFP transgene marker. To ensure the lack of GFP detection (Fig. 3b) was not due to transgene silencing, we isolated genomic DNA from more than 1000 male STSS flies and screened for presence of the transgene by PCR. We observed a clear band of the expected size from a positive control (gDNA isolated from 10 DmXL^tTA^ flies) but not in gDNA isolated from the putative non-transgenic males (Fig. 3c). Spiking trace amounts of positive control gDNA confirmed that limit of detection via this assay at less than 1:3000 transgenic:non-transgenic gDNA. This confirmed that the assay was sufficiently powerful to detect any transgenic flies that would have been present in the screened population.

## Discussion

This study demonstrated the proof of concept for STSS using *D. melanogaster* as a model system. The basic genetic parts have been demonstrated in numerous pest insects and we believe that STSS can be readily adapted to improve SIT programs by enabling efficient sex-sorting for male only release. We show here that STSS can be implemented with a variety of lethal genetic constructs, including female-lethal constructs, provided they are localized to the appropriate sex chromosome (**Supplementary Note 1**). Although we used ΦC31 mediated integration, recent advancements with CRISPR systems have found that it’s possible to target integration to many genomic loci in insects, including the repeat rich Y chromosome^20^. We did not observe any transgenic flies that were able to reproduce in the absence of tetracycline. However, the scale up of our experiments only allow us to state that the transgenic fly escape is less than 00.1% (Fig. 3c). SIT programs generate millions of animals for release on a weekly basis and it is possible that some small number of transgenic animals would be released via the STSS method, although they are likely to suffer fitness defects. USDA Organic certification in the United States prevents organic farms from utilizing GMO technology, but does not establish a detection threshold. The Non-GMO Project, a leading certifier of non-GMO status, establishes transgene threshold levels of 0.25% to 1.5% depending on the product category^21^. Thus, our application of STSS in *D. melanogaster* produces males that could in principle be certified as non-GMO.

Production of males incapable of reproduction with wild females does not necessarily require radiation treatment. The incompatible insect technique (*i.e.* IIT) relies on males infected with certain strains of *Wolbachia* bacteria. Mating between infected males and uninfected females results in embryonic lethality since the female produced egg does not contain an antidote to a deubiquitylating enzyme toxin delivered by the sperm^22^. Transgenic approaches have been developed or proposed to generate males that cannot reproduce with wild females^23–26^.

Combining STSS with cytoplasmic incompatibility would eliminate the need for any additional sterilization treatments and could enable the release of eggs/larvae to further reduce costs. Shipping of eggs, particularly of species such as *Ae. aegypti* which can be stored dry for extended periods, would allow for rearing facilities to be located far from control sites and reduce local infrastructure costs. However, this would require utilizing tetracycline resistant *Wolbachia* or non-antibiotic control of a lethal circuit by using heat-shock or some other small molecule regulators.

## Materials and Methods

### Plasmid construction

The tetracycline repressible lethal circuit was made by adapting a previously described female-lethal piggybac vector, pB[FL3]^12^. We replaced the female specific intron and 5’-UTR with a myosin heavy chain intron and syn21 translational enhancers^19^. The final plasmid, pMM7-10-1, was made by transferring the lethal circuit to pUB-EGFP^27^ which contains an *attB* site for ΦC31 mediated integration and ubiquitin promoter driven EGFP expression.

The tetracycline female lethal circuit was made by adapting a previously described female-lethal piggybac vector, pB[FL3]^16^. The final plasmid, pMM7-8-1, was made by transferring the lethal circuit to pUB-EGFP^27^, which contains an *attB* site for ΦC31 mediated integration and ubiquitin promoter driven EGFP expression. Annotated plasmid sequences for pMM7-10-1 and pMM7-8-1 are available in supplementary file STSSplasmids.zip and have been deposited in GenBank with accession numbers MN630870and MN630871, respectively.

### Generating and maintaining transgenic drosophila strains

*D. melanogaster* strains were maintained at 25°C and 12 h days in cornmeal agar (NutriFly®, Genesee Scientific, San Diego, CA) supplemented with 10-200 µg/ml tetracycline, as necessary. Transgenic repressible lethal *D. melanogaster* strains where generated by microinjection (Bestgene Inc, Ca) and ΦC31 mediated integration of pMM7-10-1 into the X-chromosome *attP* site of y[1] w[*] P{y[+t7.7]=CaryIP}su(Hw)attP8 (BDSC #32233)^28^ to make DmXL^tTA^ and the Y-chromosome *attP* of y1 w*/Dp(2;Y)G, P{CaryP}attPY^29^ to make DmYL^tTA^. Transgenic female lethal strains were generated by microinjection and ΦC31 integration of pMM7-8-1 into the X-chromosome at *attP* sites y[1] w[*] P{y[+t7.7]=CaryIP}su(Hw)attP8 (BDSC #32233)^28^ to create DMXFL1^tTA^ or y[1]w[1118]pBac{y[+]-attP-9A}VK00006 (BDSC # 9726)^30^ to create DMXFL2^tTA^. Transgenic animals were isolated by crossing to FM6 balancers, then homozygosed by selecting non-balancer animals to generate true breeding strain. DMXFL12^tTA^ flies were created by isolating recombinant chromosome of DMXFL1^tTA^ and DMXFL2^tTA^ and screening for the presence of transgene in both locations of the X-chromosome.

### Fly viability assays

Desired number of male and virgin female flies were moved to new tubes containing media either in presence or absence of tetracycline and allowed to lay eggs for five days at 25°C and 12 h light protocol. After five days, adults were removed from the tubes and offspring were allowed to develop in the incubator. Adult flies were counted as they emerged from the pupae for a total of 15 days from the start of experiment.

### PCR Verification

Fly genomic DNA was isolated in a pool by grinding in 25 µl of “Squish Buffer” (10 mM Tris, 1 mM EDTA, 25 mM NaCl, 8 U/ml ProK (NEB P8107S)) per adult. ProK was heat inactivated at 98 °C for 4 min. For transgene PCR screen, 1181 STSS males were pooled together as one sample and compared to 5 male and 5 female DMXL^tTA^ flies in a separate pool of genomic DNA as positive control. The positive control samples were diluted in with STSS gDNA in the following ratios: 1:5, 1:25, 1:125, 1:625, and 1:3125. For each reaction, 1 μL of template gDNA was used in a 20 μL PCR reaction with primers that anneal within the transgene. The following primers were used for amplification of the transgene, Fwd: 5’-GCCGCAGAATTCTCTCTATC-3’ and Rev: 5’-CTTAGCTTTCGCTTAGCGACG-3’.

## Supplementary Note 1

### X-linked female lethal constructs are also suitable for STSS

An alternative approach is to create a strain containing X-linked female lethal circuit (XFL-strain, Supplementary Fig. 1a), which when active is selectively lethal to females containing the circuit. Alternative splicing of the tTA transgene based on the sex-determining *tra* gene from *C. hominivorax*^11^ yielded the male sex selection (Supplementary Fig. 1b). Males produced from this strain in the selection media when crossed with females obtained from YL strain would result in non-transgenic males (Supplementary Fig. 1c). We found a dosage-dependent phenotype for the female lethal construct, with two copies placed at distinct loci on the X-chromosome required for lethality in hemizygous female offspring (Supplementary Fig. 1d,e). When the two-stage mating scheme was implemented, we observed a strong enrichment for non-transgenic males (Supplementary Fig. 1f), however it was not as strong as for the repressible-lethal system described above. Most experimental replicates produced a small number of females that were GFP+, although all appeared sickly. Although it is possible that these females are fertile, they often died on the vial food within 3 days of eclosion, highlighting the effectiveness of tTa transactivator. These data highlight the fact either bi-sex lethal or female lethal genetic constructs can be used for STSS as long as they are localized to the appropriate sex chromosomes.

**Supplemental Fig. 1.**
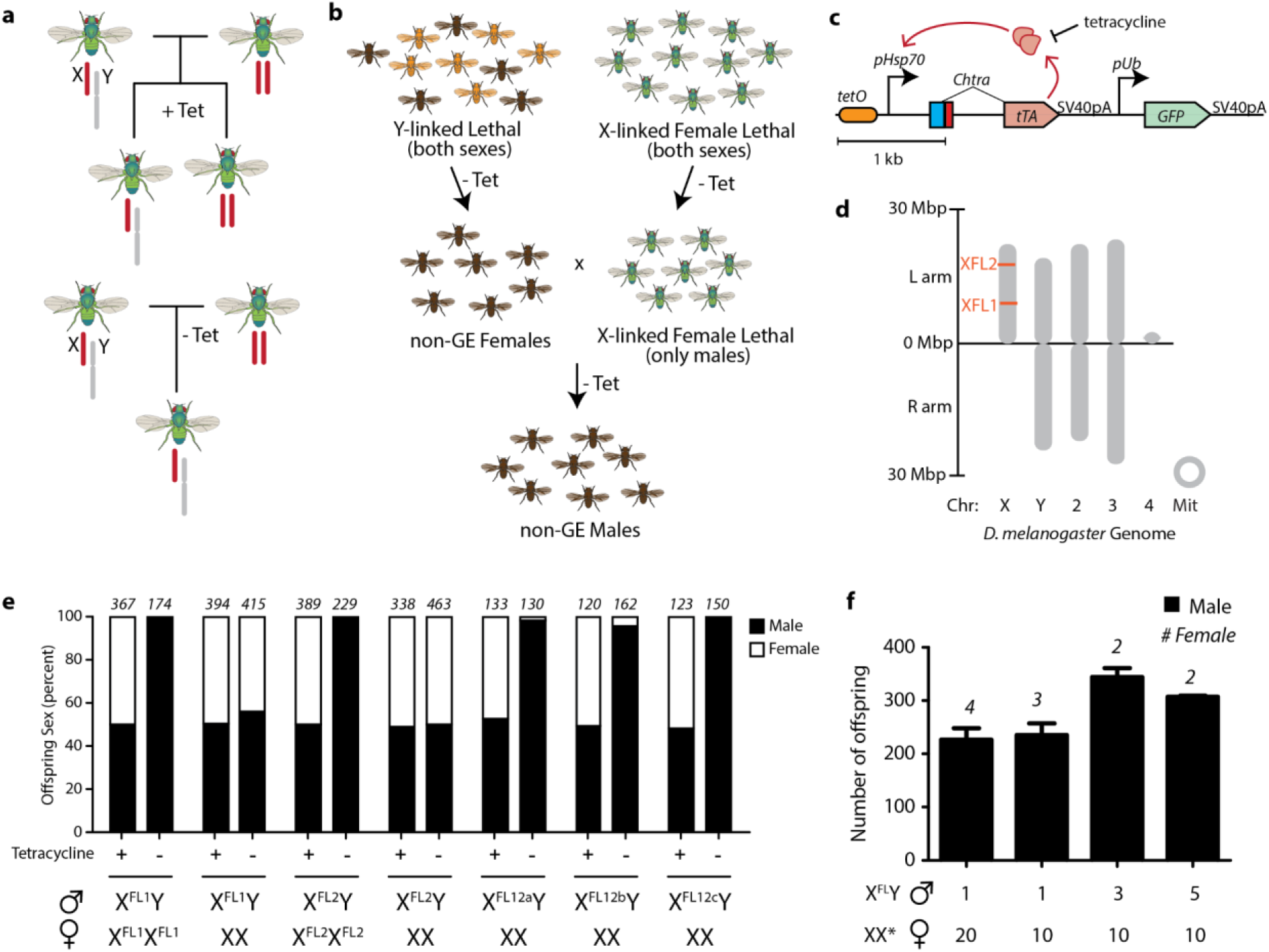
An alternative approach of STSS. (**a**) Reproductive behaviour of X-linked Female Lethal construct used in female-lethal (FL-) STSS. (**b**) Mating scheme for producing non-transgenic males via FL-STSS. Combining non-transgenic females produced from the YL strain with adult male flies produced from the XFL strain results in death of all offspring except for non-transgenic males. (**c**) Genetic design of FL construct. (**d**) Chromosomal location of FL constructs in two copy ‘FL12a-c’ flies. FL1 and FL2 have only one copy of the X-linked FL construct on their X-chromosome. (**e**) Proportion of male and female offspring generated from self-mating or outcrosses to non-transgenic (w1118) for DmXFL1, DmXFL2, and three independently generated DmXFL12 genotypes. Parental genotypes are indicated below the x-axis. Results are shown in the presence or absence of tetracycline. Numbers above bars indicate total number of progeny produced from at least three biological replicates. (**f**) Average number of adult males obtained from mating between different proportions of non-transgenic female flies obtained from DmYL^tTA^ in absence of tetracycline (‘XX*’) when combined with adult male DmXFL12c flies in absence of tetracycline. Data represent mean numbers from 2 biological replicates with error bars showing standard error of the mean (s.e.m.). Average numbers of females produced are indicated numerically above bars. Total numbers from all replicates are (from left to right): N=460, N=475, N=692, N=617. Numbers below x-axis indicate number of parental flies of each genotype.

## References

1. Dyck, V. A., Hendrichs, J. & Robinson, A. S. Sterile insect technique: principles and practice in area-wide integrated pest management. (Springer, 2005).

2. Wyss, J. H. Screwworm eradication in the Americas. Ann. N. Y. Acad. Sci. 916, 186–93 (2000).

3. Vargas-Terán, M., Hofmann, H. C. & Tweddle, N. E. Impact of Screwworm Eradication Programmes Using the Sterile Insect Technique. in Sterile Insect Technique 629–650 (Springer-Verlag, 2005). doi:10.1007/1-4020-4051-2_24

4. Hendrichs, J., Robinson, A. S., Cayol, J. P. & Enkerlin, W. Medfly areawide sterile insect technique programmes for suppression or eradication: The importance of mating behavior studies. Florida Entomol. 85, 1–13 (2002).

5. Dominiak, B. C. et al. Evaluating irradiation dose for sterility induction and quality control of mass-produced fruit fly Bactrocera tryoni (Diptera: Tephritidae). J. Econ. Entomol. 107, 1172–8 (2014).

6. Rendón, P., McInnis, D., Lance, D. & Stewart, J. Medfly (Diptera:Tephritidae) Genetic Sexing: Large-Scale Field Comparison of Males-Only and Bisexual Sterile Fly Releases in Guatemala. J. Econ. Entomol. 97, 1547–1553 (2004).

7. Carvalho, D. O. et al. Mass Production of Genetically Modified Aedes aegypti for Field Releases in Brazil. J. Vis. Exp. e3579 (2014). doi:10.3791/3579

8. Marois, E. et al. High-throughput sorting of mosquito larvae for laboratory studies and for future vector control interventions. Malar. J. 11, 302 (2012).

9. Caceres, C. Mass rearing of temperature sensitive genetic sexing strains in the Mediterranean fruit fly (Ceratitis capitata). Genetica 116, 107–16 (2002).

10. Fu, G. et al. Female-specific flightless phenotype for mosquito control. Proc. Natl. Acad. Sci. 107, 4550–4554 (2010).

11. Concha, C. et al. A transgenic male-only strain of the New World screwworm for an improved control program using the sterile insect technique. BMC Biol. 14, (2016).

12. Yan, Y., Linger, R. J. & Scott, M. J. Building early-larval sexing systems for genetic control of the Australian sheep blow fly Lucilia cuprina using two constitutive promoters. Sci. Rep. 7, 2538 (2017).

13. Jin, L. et al. Engineered female-specific lethality for control of pest lepidoptera. ACS Synth. Biol. 2, 160–166 (2013).

14. Tan, A. et al. Transgene-based, female-specific lethality system for genetic sexing of the silkworm, Bombyx mori. Proc. Natl. Acad. Sci. U. S. A. 110, 6766–70 (2013).

15. Smanski, M. J. & Zarkower, D. Genetic manipulation of sex ratio in mammals: the Reaper comes for Mickey. EMBO Rep. 20, (2019).

16. Li, F., Wantuch, H. A., Linger, R. J., Belikoff, E. J. & Scott, M. J. Transgenic sexing system for genetic control of the Australian sheep blow fly Lucilia cuprina. Insect Biochem. Mol. Biol. 51, 80–88 (2014).

17. Heinrich, J. C. & Scott, M. J. A repressible female-specific lethal genetic system for making transgenic insect strains suitable for a sterile-release program. Proc. Natl. Acad. Sci. U. S. A. 97, 8229–32 (2000).

18. Fu, G. et al. Female-specific insect lethality engineered using alternative splicing. Nat. Biotechnol. 25, (2007).

19. Pfeiffer, B. D., Truman, J. W. & Rubin, G. M. Using translational enhancers to increase transgene expression in Drosophila. Proc. Natl. Acad. Sci. U. S. A. 109, 6626–31 (2012).

20. Buchman, A. & Akbari, O. S. Site-specific transgenesis of the Drosophila melanogaster Y-chromosome using CRISPR/Cas9. Insect Mol. Biol. 28, 65–73 (2019).

21. Bain, C. & Selfa, T. Non-GMO vs organic labels: purity or process guarantees in a GMO contaminated landscape. Agric. Human Values 34, 805–818 (2017).

22. Beckmann, J. F., Ronau, J. A. & Hochstrasser, M. A Wolbachia deubiquitylating enzyme induces cytoplasmic incompatibility. Nat. Microbiol. 2, 17007 (2017).

23. Kandul, N. P. et al. Transforming insect population control with precision guided sterile males with demonstration in flies. Nat. Commun. 10, 84 (2019).

24. Maselko, M., Heinsch, S. C., Chacón, J. M., Harcombe, W. R. & Smanski, M. J. Engineering species-like barriers to sexual reproduction. doi:10.1038/s41467-017-01007-3

25. Waters, A. J. et al. Rationally-engineered reproductive barriers using CRISPR & CRISPRa: an evaluation of the synthetic species concept in Drosophila melanogaster. Sci. Transl. Med. (2018). doi:10.1101/259010

26. Thomas, D. D., Donnelly, C. A., Wood, R. J. & Alphey, L. S. Insect population control using a dominant, repressible, lethal genetic system. Science (80-.). 287, 2474–2476 (2000).

27. Schetelig, M. F. et al. Site-specific recombination for the modification of transgenic strains of the Mediterranean fruit fly Ceratitis capitata. Proc. Natl. Acad. Sci. U. S. A. 106, 18171–6 (2009).

28. Pfeiffer, B. D. et al. Refinement of Tools for Targeted Gene Expression in Drosophila. Genetics 186, 735–755 (2010).

29. Szabad, J., Bellen, H. J. & Venken, K. J. T. An assay to detect in vivo Y chromosome loss in Drosophila wing disc cells. G3 Genes, Genomes, Genet. 2, 1095–1102 (2012).

30. Venken, K. J. T., He, Y., Hoskins, R. A. & Bellen, H. J. P[acman]: A BAC transgenic platform for targeted insertion of large DNA fragments in D. melanogaster. Science (80-.). 314, 1747–1751 (2006).

